# Specialized response of default mode subnetworks and multiple-demand regions to transitions of person, place and time

**DOI:** 10.1101/2025.01.14.631412

**Authors:** Ashley X Zhou, John Duncan, Daniel J Mitchell

**Affiliations:** MRC Cognition and Brain Sciences Unit, University of Cambridge

**Keywords:** functional MRI, default mode network, cognition, naturalistic stimuli, transitions

## Abstract

This study used functional MRI data from the *StudyForrest* dataset to investigate the role of subnetworks of the default mode network (DMN) during naturalistic stimulus transitions of different types and magnitudes. We found distinct activation profiles within the DMN: the dmPFC subnetwork was specifically associated with character and location transitions, the MTL subnetwork preferred location and temporal transitions, while the Core DMN subnetwork responded to all three transition types. The multiple-demand network instead responded selectively to temporal transitions. These distinct response profiles appeared largely invariant to the semantic distance implied by the transitions. All subnetworks also responded significantly, and in a graded manner, to subjective event boundaries. Results suggest specific roles of the DMN subnetworks in perceiving and segmenting naturalistic events, supporting the view that DMN subnetworks cooperate in interpreting continuous external events and maintaining an updated contextual model of the world.

## Introduction

Since the early findings of Shulman et al. (1997) and Raichle et al. (2001), many studies have shown deactivation of the default mode network (DMN) during external tasks compared to rest. On this basis the DMN has often been considered a ‘task-negative’ network (Fox et al., 2005). Subsequent studies have refined this view (Spreng, 2012), finding DMN activation associated with specific aspects of internally-oriented cognition (Buckner et al., 2008), such as self-related thought (Andrews-Hanna et al., 2010; Davey et al., 2016; Gusnard et al., 2001), social cognition (Mars et al., 2012), autobiographical memory retrieval and mental imagery (Addis et al., 2007; Buckner & Carroll, 2007; Hassabis & Maguire, 2009; Schacter et al., 2007). Recent studies have also identified Core DMN activity during externally oriented cognition, including automatized responding (Vatansever et al., 2017) as well as more effortful instances of cognitive control such as at visually-cued switches between multiple tasks, compared to task repeats (Crittenden et al., 2015; Kurtin et al., 2023; Smith et al., 2018; Zhou et al., 2024b).

Of the various theories of DMN function, one that can accommodate observed activations at external task transitions as well as during restful and introspective thought is the ‘sentinel hypothesis’. This early proposal summarised the DMN’s role as serving to monitor the internal and external environment, facilitating adaptive responses to changes in current conditions (Gilbert et al., 2007; Raichle et al., 2001). Relatedly, the DMN and associated networks have been hypothesised to process ‘situation models’ that represent the setting in which behaviour occurs (Ranganath & Ritchey, 2012). The suggestion is that the DMN is involved in monitoring and updating a mental model, including representation of conditions and features, both external and internal, which provide broad context to events and behaviour, whether spontaneously generated during rest, or in the service of an imposed task.

The DMN’s potential role in representing or updating such a mental model, particularly in response to external stimulus changes, may have eluded traditional task-based paradigms that emphasize isolated focal tasks without reflecting a more complex context. To address this limitation and investigate network activity in response to richer and more dynamic cognitive demands, recent literature has increasingly moved towards the use of continuous and more naturalistic stimuli that demand real-time information integration, often from multiple modalities (Menon, 2023; Sonkusare et al., 2019). By integrating extrinsic and intrinsic information during narrative processing, these studies provide a closer resemblance to real-world processing than more highly controlled tasks (Bottenhorn et al., 2018). Widespread brain regions are known to respond to narrative transitions of many kinds (Speer et al., 2009; Zacks et al., 2010). Interestingly, a parametric response to multiple transitions overlaps with default mode regions, especially in the medial parietal lobe (Speer et al., 2009). However, it is unclear how these activations relate to DMN subnetworks, and whether they depend on the semantic magnitude of the underlying transitions.

Within the DMN, distinct subnetworks are now thought to be relatively specialized for different cognitive processes (Andrews-Hanna et al., 2010; Andrews-Hanna, 2012; Andrews-Hanna et al., 2014; Axelrod et al., 2017; Wen, Mitchell, et al., 2020). A medial temporal lobe (MTL) subnetwork (also including ventromedial prefrontal cortex, retrosplenial cortex and the posterior inferior parietal lobule) is particularly implicated in memory-based scene construction (Hassabis et al., 2007; Palombo et al., 2018; Ranganath & Ritchey, 2012), and overlaps with medial temporal lobe and retrosplenial brain regions that are activated during spatial navigation tasks (Epstein, 2008). Zhou et al. (2024a) found that activation patterns within the DMN’s MTL subnetwork represented the category of background scenes, further suggesting a role in processing contextual location information. A dorsomedial prefrontal cortex (dMPFC) subnetwork (also including the temporoparietal junction and anterior and lateral temporal cortex) has been particularly associated with social cognition, including theory of mind and personality judgements (Denny et al., 2012; Gallagher et al., 2000; Hassabis et al., 2014; Saxe & Kanwisher, 2003; Tavares et al., 2008; Van Overwalle, 2009), although it is also implicated in semantic and conceptual cognition more generally (Humphreys et al., 2015). Finally, a midline Core subnetwork (including posterior cingulate and anterior medial prefrontal regions) may integrate information from the other two subsystems from a self-referential perspective (Johnson et al., 2002; Kelley et al., 2002; Kurczek et al., 2015; Qin & Northoff, 2011; Sajonz et al., 2010). Overall, the literature suggests that the DMN subnetworks may respond to different aspects of a mental model, including social and spatial features, and might support their integration into a coherent mental model of the world to guide behavior and decision-making.

Given the DMN’s proposed role in monitoring changes to a broadly-defined mental model, changes in features contributing to the current model are expected to elicit DMN activation, consistent with its response at cognitive task switches (Crittenden et al., 2015; Smith et al., 2018; Zhou et al., 2024b, 2024a), and narrative transitions (Speer et al., 2009; Yazin et al., 2024; Zacks et al., 2010), and with changes in activity pattern at narrative event boundaries (Baldassano et al., 2017). This study further investigates a role of the DMN in representing and updating a mental model, by comparing recruitment of its subnetworks at various types and magnitudes of transitions in naturalistic stimuli. Specifically, we first aimed to identify distinct events that differentially drive activation of DMN subnetworks as participants watch a movie. A first exploratory analysis used the movie-watching fMRI data from the Human Connectome Project (HCP) (Van Essen et al., 2013) to generate hypotheses regarding transition types that differentially activate DMN subnetworks. A second set of analyses then assessed and extended these hypotheses using fMRI data from the *StudyForrest* project (Hanke et al., 2016), by examining how the DMN subnetworks responded to different types and magnitudes of narrative transitions, as well as at subjective event boundaries. The DMN’s response to narrative transitions was also compared to the response of the multiple demand (MD) network (Duncan, 2010), widely implicated in cognitive control. While much previous work has characterized MD activity as ‘task-positive’, and often anti-correlated with the DMN (Fox et al., 2005), the two networks can co-activate at task transitions compared to task repeats (Zhou et al., 2024a) and at boundaries between natural task sequences (Wen, Duncan, et al., 2020).

### Hypothesis generation using the HCP movie-watching dataset

#### Participants

The Human Connectome Project (HCP) movie-watching dataset (Van Essen et al., 2013) included fMRI data from 184 participants (94 female), all of whom were healthy young adults recruited from the general population. Participants were aged 22–35, with a mean age of 29.4 years. All participants were screened for neurological and psychiatric conditions. The study was approved by the Washington University in St. Louis Institutional Review Board (IRB).

#### MRI data acquisition and preprocessing

Exploratory analyses were conducted using the HCP movie-watching fMRI data from the 7T release (https://www.humanconnectome.org/study/hcp-young-adult). Participants viewed four runs of short film clips while fMRI data were acquired with a 7 Tesla Siemens Magnetom scanner at the University of Minnesota Center for Magnetic Resonance Research. We used data from the four movie-watching runs, acquired across two sessions, with 177-182 participants’ data available per run. Movie runs were acquired using gradient echo-planar imaging (EPI) (TR = 1000ms, TE= 22.2 ms, flip angle 45°, voxel size 1.6 mm^3^, FOV= 208 x 208 mm, 85 axial slices, multiband factor = 5, partial Fourier sampling = 7/8, imaging acceleration factor (iPAT) = 2, echo spacing =0.64 ms, bandwidth = 1924 Hz/Px) (Finn & Bandettini, 2021). Phase encoding direction alternated between posterior to anterior (PA) for movie runs 2 and 3, and anterior to posterior (AP) for movie runs 1 and 4. Each of the four movie-watching runs (15.0-15.3 minutes) combined four or five short clips from independent and Hollywood movies ranging from 1 to 4.3 minutes in length. At the start and end of each run, and between each clip, a black screen was presented for 20 s, with the instruction “REST”. Eye movements were monitored but not controlled. The last clip of each run contained the same montage of brief video snippets for validation analyses.

Functional images of the movie runs had been preprocessed using the HCP pipeline (Glasser et al., 2013), which included spatial distortion correction, motion correction, registration of functional to structural scans, non-linear registration to MNI space and grand-mean intensity normalization. Data were smoothed by a 2mm kernel, and temporally filtered with a Gaussian-weighted linear high-pass filter with a cutoff of 200s. ICA+FIX (Salimi-Khorshidi et al., 2014) was used to reduce spatially specific noise. Functional images had been mapped from volume to surface space using ribbon-constrained mapping, and a multimodal surface matching algorithm (MSMAll; Robinson et al., 2014) for alignment of cortical areas. For each movie run, we z-scored fMRI activity over time, per vertex, and averaged across all subjects (n=177-182).

#### Regions of Interest (ROIs)

Surface-based ROIs for the three DMN subnetworks were taken from the Yeo 17 parcellation (Yeo et al., 2011, network numbers 15, 16 and 17; which correspond to the “Core”, “MTL” and “dMPFC” subnetworks proposed by Andrews-Hanna et al., 2014). A surface-based ROI for the core MD network was taken from Assem et al. (2020). Data were averaged across vertices within each ROI.

#### Analysis

Peaks and troughs of each DMN subnetwork’s activity were examined, focusing on identifying common features that the DMN subnetworks were most attuned to. For each subnetwork, the top ten peaks and troughs were selected, with minimum separation of 10 seconds between the local peaks, and excluding peaks within the “rest” periods. To account for the hemodynamic response, scenes 4, 6 and 8 seconds before each peak were identified. Then, candidate features, consisting of long shots of scenes, transitions of scenes, shots including characters, and transitions of characters, in the scenes preceding peaks and troughs, were counted manually by the experimenter, as they were noticed to be prominent features within the selected frames. Only characters that were discernable and distinct were counted (for example, silhouettes in the distance and close-up shots of body parts were not counted). Any type of transition to a character was counted, including from long shots of scenes to a character, or from one character to another. Similarly, transitions to scenes included transitioning from a close-up shot or character to the surrounding scene, as well as transitions from one scene to another. Note that transitions to new characters and to new scenes were also included in the simple count of frames including characters and scenes.

## Results

Responses from the three DMN subnetworks were plotted across the time course of each movie, and movie frames corresponding to 4, 6 and 8 seconds before the ten largest peaks and troughs were plotted alongside, to show the corresponding events on the screen while accounting for the delayed hemodynamic response.

These movie frames were then investigated and compared for each movie and each ROI. Table 1 describes the distribution of the counted features: the presence of characters and long shots of scenes, and transitions to characters and to long shots of scenes. For each feature and each DMN subnetwork, the table show the range of the number of peak frames containing these features, out of ten per run, and the overall percentage of peaks showing these features, out of all 40 peaks.

**Table 1.** Description of the frames preceding the top 10 activation peaks and troughs across the four runs, for each DMN subnetwork ROI. The numbers and percentages of frames containing characters, scenes, and transitions of these features are reported, for the frames 4, 6, and 8 seconds before each activity peak/trough.

| dmPFC |  |  |  |  |
| --- | --- | --- | --- | --- |
| Description | Peaks per run | % of peaks | Troughs per run | % of troughs |
| Characters/Faces | 7-10 | 85.0 | 2-5 | 37.5 |
| Transition of characters | 3-10 | 62.5 | 1-5 | 20.0 |
| Long shot of scenes | 0-3 | 12.5 | 1-5 | 35.0 |
| Transition of scenes | 0-2 | 10.0 | 0-4 | 15.0 |

| MTL |  |  |  |  |
| --- | --- | --- | --- | --- |
| Description | Peaks per run | % of peaks | Troughs per run | % of troughs |
| Characters/Faces | 1-8 | 50.0 | 5-10 | 77.5 |
| Transition of characters | 0-4 | 15.0 | 1-2 | 10.0 |
| Long shot of scenes | 6-10 | 82.5 | 0-3 | 7.5 |
| Transition of scenes | 7-9 | 77.5 | 0-3 | 7.5 |

**Table 1.** Description of the frames preceding the top 10 activation peaks and troughs across the four runs, for each DMN subnetwork ROI.
| Core DMN |  |  |  |  |
| --- | --- | --- | --- | --- |
| Description | Peaks per run | % of peaks | Troughs per run | % of troughs |
| Characters/Faces | 5-10 | 77.5 | 3-8 | 47.5 |
| Transition of characters | 3-9 | 55.0 | 0-5 | 20.5 |
| Long shot of scenes | 2-6 | 37.5 | 2-6 | 35.0 |
| Transition of scenes | 2-4 | 27.5 | 1-2 | 12.5 |

First, it was noted that for the dmPFC subnetwork, the frames corresponding to activity peaks consistently, and almost exclusively, showed faces or characters, with 85% of frames at peak activity showing characters. This was in stark contrast to the frames from the peaks of MTL subnetwork activity, which primarily displayed medium to long shots (82.5%), which in film refers to shots taken where the character does not take up much of the frame and that primarily show where the scene takes place. Core DMN’s activity peaks corresponded to a mixed set of features including both characters (77.5%) and scenes (37.5%).

Next, frames 4, 6 and 8 seconds before the activity peaks were manually counted for times when they contained a transition to a new character or scene. The number and proportion of top peak frames for the subnetworks, recorded in the table, show a clear pattern that suggests a preference for character transitions for the dmPFC subnetwork (62.5% of peaks, corresponding to 73.5% of the peak frames including characters), a preference for scene transitions for the MTL subnetwork (77.5% of peaks, corresponding to 93.9% of the peak frames including scenes), and a mixture for the Core subnetwork, often including transitions of character (55% of peaks; 71.0% of peak frames including characters) or scene (27.5% of peaks; 73.3% of peak frames including scenes).

For each subnetwork, we also counted the number of features in the frames immediately preceding the activity troughs. The opposite pattern was observed. Most (77.5%) MTL subnetwork activity troughs corresponded to the presence of characters on the screen, while only 7.5% of troughs contained scenes or transitions of scenes. Conversely, only 20% of troughs for dmPFC subnetwork activity showed transition of characters, further supporting the hypothesis that character transitions drive dmPFC subnetwork activity, and scene transitions drive MTL subnetwork activity, and that the two networks show differentiated behaviour in response to these features of the naturalistic stimuli.

## Discussion

The exploratory analyses of the HCP movies dataset suggested that the DMN subnetworks responded differently to movie events. The dmPFC subnetwork showed a preference for characters and especially for character transitions, the MTL subnetwork preferred scenes and especially scene transitions, and the Core subnetwork responded to a mixture of both. These findings are broadly in line with previous literature, implicating the dmPFC subnetwork especially in social cognition, and the MTL subnetwork in spatial cognition (Andrews-Hanna et al., 2010, 2014; Wen, Mitchell, et al., 2020).

These observations were not analysed with formal statistical tests due to implicit consideration of multiple alternative features. Instead, they formed the basis of predictions to be tested in a separate dataset. Based on these observations, the following hypotheses were proposed for further testing. First, increased activity of dmPFC, MTL and Core subnetworks of the DMN was hypothesised in response to character transitions, location transitions and both transitions respectively. Additionally, considering that many of these transitions occur concurrently with temporal shifts in the narrative, we hypothesized that the subnetworks might also respond to temporal transitions.

Since we have previously observed Core DMN and MD regions to co-activate at transitions between tasks (Smith et al., 2018; Wen, Duncan, et al., 2020; Zhou et al., 2024a), and MD activity is also observed at musical transitions (Sridharan et al., 2007), it was hypothesised that MD regions might also respond to narrative transitions, although we were agnostic regarding any potential feature preference.

To test these hypotheses, and further explore the role of DMN in context monitoring, a different dataset, *StudyForrest* (Hanke et al., 2014), was used in the second set of analyses. This dataset includes annotations that can be used to isolate the specific feature transitions of interest (character, location and temporal transitions), and to test for their effect on the DMN subnetworks. Furthermore, the dataset has been expanded to include subjective annotations of event boundaries, where multiple observers indicated meaningful transitions within the narrative (Ben-Yakov & Henson, 2018). The degree of boundary “salience” was determined by the convergence of observer annotations, providing a measure of the perceived significance of each event boundary. Therefore, we also assessed how DMN activity extends to subjectively salient narrative transitions.

In previous studies, the Core DMN has shown stronger responses to task transitions of greater magnitude (within versus between stimulus domains) (Crittenden et al., 2015; Smith et al., 2018; Zhou et al., 2024b). To test whether DMN subnetwork activity also depends on the semantic magnitude of naturalistic transitions. we also defined and compared different levels of magnitude for each transition type.

## Hypothesis testing using the StudyForrest dataset

### Methods

#### Participants

The *StudyForrest’s* two hour audio-visual dataset included fMRI data from 15 right-handed participants (10 female), all of whom were native German speakers, who volunteered to view the German-dubbed version of the film “Forrest Gump” (R. Zemeckis, Paramount Pictures, 1994) during MRI scanning (Hanke et al., 2016; Sengupta et al., 2016). Participants were aged 21-39, with a mean age of 29.4 years. All had normal vision, assessed at the Visual Processing Laboratory, Ophthalmic Department in Otto-von-Guericke University. Participants provided informed consent, and received monetary compensation. The study was approved by the Otto-von-Guericke University’s Ethics Committee. All participants were familiar with the film, as they had previously heard an audio-only version in another study (Hanke et al., 2014), and all except one indicated that they had previously seen the film.

#### MRI data acquisition and pre-processing

The German-dubbed film of “Forrest Gump” had been edited down to two hours, and segmented into eight runs of 11-18 minutes (Hanke et al., 2016). Data were acquired on a 3 Tesla Philips Achieva dStream MRI Scanner (Philips Medical Systems) with a 32-channel head coil (Hanke et al., 2016). During each run, functional volumes were acquired using a gradient-echo, T2*-weighted EPI sequence (TR = 2000ms, TE= 30ms, flip angle = 90°, voxel size = 3 x 3 x 3 mm, field of view = 240 x 240 mm, 35 axial slices acquired in ascending order with a 10% gap). T1-weighted structural scans were acquired using a 3D turbo field echo sequence (TR_= 2500 ms, TE = 5.7 ms, TI = 900 ms, flip angle = 8°, field of view = 191.8_x 256_x 256 mm, with 0.67 mm isotropic resolution). Further imaging details are given in Hanke et al.(2014, 2016).

Preprocessing was performed by Ben-Yakov and Henson (2018) in Matlab (The Mathwords Inc), using the automatic analysis toolbox (Cusack et al., 2015) to call SPM12 functions (Wellcome Department of Cognitive Neurology, London, UK). Pre-processing steps included spatial realignment of functional volumes, slice-time correction by interpolating to the middle slice, rigid body co-registration of functional data to the T1 anatomical images, nonlinear spatial normalisation to the Montreal Neurological Institute (MNI) template, and reducing effects of abrupt motion by applying wavelet despiking (Ben-Yakov & Henson, 2018).

#### Regions of Interest (ROIs)

Primary analyses focus on the subnetworks of the default mode network (Core, MTL and dmPFC), with volumetric ROIs taken from Wen, Mitchell, et al. (2020). The Core subnetwork included the anterior medial prefrontal cortex, and the posterior cingulate cortex. The MTL subnetwork included the retrosplenial cortex, parahippocampal cortex, posterior inferior parietal lobule, hippocampal formation, and ventral medial prefrontal cortex. The dmPFC subnetwork included the dorsomedial prefrontal cortex, the temporal pole, the temporoparietal junction, and lateral temporal cortex (Andrews-Hanna, 2012). ROIs for these subnetworks were based on networks 10, 15, 16 and 17 from the Yeo et al (2011) 17-network parcellation, subdivided according to the coordinates provided in Andrews-Hanna et al. (2012) for each of the three subnetworks (see Wen, Mitchell, et al., 2020). A volumetric ROI for the frontoparietal multiple demand network was taken from Mitchell et al. (2016). For the analysis of subjective event boundaries, bilateral group ROIs of the hippocampus had been traced by (Ben-Yakov & Henson, 2018), and are included here to reproduce their results.

#### Subjective event boundaries

Subjective annotations from Ben-Yakov & Henson were used to identify narrative event boundaries (Ben-Yakov & Henson, 2018). Ben-Yakov & Henson’s event boundaries were collected from 16 participants who were independent from the cohort that viewed the film in the scanner, and were asked to indicate, by pressing a key, while watching the film, “when they felt one event (meaningful unit) ended and another began” in the manner that felt most natural. Eight participants watched the German-dubbed version and eight watched the English version of the film. 0.9 s were subtracted from button presses based on estimated reaction times during prior testing. Marked boundaries were combined if different participants marked them within 2 TRs (2s per TR). Finally, boundaries that were identified by fewer than five participants were excluded from further analysis. See Ben-Yakov and Henson (2018) for further details.

There were a total of 161 boundaries, with 12-25 per run, included in the analysis. Boundaries were assigned to one of three “salience” levels, based on the degree of agreement between participants, and with a similar number of boundaries per level. Low salience event boundaries were marked by 5-6 participants, medium salience event boundaries were marked by 7-9 participants, and high salience event boundaries were marked by 10 or more participants. For the low, medium and high salience levels there were 60, 43, and 54 event boundaries respectively. The subgroups that watched the German-dubbed and English versions had no significant difference in their boundary annotations.

#### Character, location, and temporal transitions

Based on the exploratory observations from the HCP dataset, additional transitions were chosen: location, temporal and character transitions. Each type of transition was then further separated into two degrees of magnitude, shown in Table 2 and explained in detail below.

**Table 2.** Distribution of different transition types across the movie runs. Transitions in location, character and time are divided into large and small changes, with descriptions of each recorded. For transitions of each type, columns show the mean and standard deviation of number of transitions per run, and the mean and standard deviation of time (s) in between each transition per run.

|  | Transition Type | Transitions in Naturalistic Stimuli (StudyForrest) |  |
| --- | --- | --- | --- |
|  |  | Number per run | Time between transitions |
| Large Location | Changes in major location and setting, e.g. different part of town | 22.50 (8.12) | 23.58 (7.49) |
| Small Location | Changes in locale, e.g. different rooms | 11.00 (11.54) | 67.97 (90.76) |
| Large Temporal | Jumps into the future from several hours to years | 16.88 (7.00) | 55.32 (23.68) |
| Small Temporal | Forward in time, e.g. small shift within one day | 7.63 (4.10) | 108.02 (83.02) |
| Large Character | First appearance of character in new scene | 35.75 (10.40) | 23.58 (7.49) |
| Small Character | Transitions between characters in a scene | 37.62 (14.11) | 23.21 (6.91) |

#### Location

In a separate study, locations within the film, and temporal progression, were manually annotated (Häusler & Hanke, 2016). The movie was inspected frame-by-frame, across four passes by the same observer, to identify and validate the annotated properties. Location was noted with three levels of detail describing the depicted scenery – Major location, Setting and Locale. While Major location identifies the location at a coarse level, such as a town, county or region, Setting distinguishes the place of the scene in a finer level, such as Gump’s house. Further, Locale details the exact area when there is more than one locale for the Setting, such as Gump’s room and Gump’s mother’s room. From the annotations of the locations and their onset times, we noted the times when the location changed from the previous frame. To assess the sensitivity to the magnitude of change from the previous scene, we combined changes in Setting or Major Location in the first group (large location changes), and put changes in Locale in the second group (small location changes).

#### Temporal progression

The flow of time in the film is dynamic, as the narrative of the film includes non-linear progressions in time, including flashbacks and substantial jumps forward in time. In the storytelling, a moment may be proceeded by an event that happened years before, or months later. To capture this, annotations of each frame noted the temporal progression based on the previous and current shot (Häusler & Hanke, 2016). The temporal progression label had four categories, namely, no discernable break in time, flashbacks (regardless of the temporal distance into the past), noticeable forward jumps in time ranging from several seconds to a few hours, and major jumps spanning from larger jumps within one day, such as change from night to day, to several years. In the following analyses, small and major forward jumps in time are used as the two levels of magnitude of temporal transition. While flashbacks were also modelled, they were not used in direct comparison to the other temporal shifts as they may be qualitatively different in nature.

#### Character transitions

In another study, nine participants had independently labeled the audio-visual movie time points with emotions of the characters (Labs et al., 2015). Here, shots were used as units for annotation, as the characters portraying emotion were slowly changing. There were 205 units across the movie stimulus and each unit was labeled with the character name and the emotion properties. In the following analyses, we used only the character label to generate a vector of time points where the character on screen changed from the previous shot. For the larger magnitude group, we used the event boundaries collapsed across all salience levels (described above) to first segment the movie, and then identified the moments where a character appeared in a segment for the first time. Transitions back to the same character within a segment formed the small magnitude group. This approach was based on the hypothesis that characters appearing for the first time in a new event may require more cognitive processing of character information compared to when the shot transitions back to a character that is already known to be present in the scene.

#### Visual and auditory confounds (z-scored)

To separate the effects of character, location and temporal transitions from shifts in lower-level perceptual features of the movie, various visual and auditory measures were also modelled as potential confounds. The confound variables below were taken from Ben-Yakov & Henson (2018), and from measures annotated by the *StudyForrest* project (Hanke et al., 2016).

*Frame_dist:* the visual difference between the current frame and the previous frame, using Image Euclidean distance (IMED). Images of the pair of frames were down-sampled by a factor of eight and then the IMED between frames was calculated, taking into consideration surrounding pixels so that the measure is less sensitive to small movements between frames.

*Frame_corr:* the spatial correlation between the current frame and previous frame, using Image Normalized Cross-Correlation (IMNCC). This uses a similar approach to IMED and the same parameters, also considering spatial relationships of pixels (Nakhmani & Tannenbaum, 2013).

*Frame_mean_lum:* the average of luminance over all pixels in each frame.

*Frame_lum_diff:* the absolute difference in average global luminance across all pixels for each pair of adjacent frames, to capture the change in global lighting.

*Rms_audio:* root-mean-square of the audio stream volume across the left and right inputs.

*Rmn_audio_diff:* the derivative of the volume function, i.e. the change in volume from the previous frame.

#### Analysis

Data for each participant were examined using a series of General Linear Models (GLM). A first GLM included separate regressors for each combination of transition magnitude (small or large), and types of transitions (location, temporal, and character transitions). Regressors were modelled by delta functions placed at the onsets of the transitions, convolved with the canonical hemodynamic response function. A second GLM was created with regressors modelled by delta functions at the low, medium and high salience onsets of subjective event boundaries (Ben-Yakov & Henson, 2018), convolved with the hemodynamic response function. For both models, collinearity between regressors was assessed using Pearson correlations and variance inflation factors (VIF).

To further isolate independent effects of each transition type, we next investigated how often transitions of different types co-occurred. We created a third GLM that used separate regressors for transitions of each type that occurred either on their own or in conjunction with transitions of another type. For this analysis, transition magnitudes were collapsed, and subjective event boundaries were considered alongside objective feature transitions. Each transition was defined as “pure” if there were no transitions of a different type within plus-or-minus two seconds. Otherwise, the transition was assigned to a separate regressor coding the particular combination of transitions that co-occurred within this time window. Therefore, the model contained 15 regressors covering all possible combinations of transitions: C, L, T, E, CL, CT, CE, LT, LE, TE, CLT, CLE, CTE, LTE, and CLTE, where C, L, T & E respectively indicate character, location and temporal transitions, and event boundaries. For this analysis, we were interested in pure transitions of each type, i.e. activation uniquely associated with regressors C, L, T and E.

For all models, the visual and audio confounds were included as parametric modulators of a single regressor with one event per TR, but are not considered further in the analyses. Six rigid-body realignment parameters were included for each run of all models, along with run means, as non-convolved regressors. A temporal high-pass filter with a default cut-off period of 128 s was applied to the data and each model.

#### Statistical Analysis

Regression coefficients from the per-participant GLMs, averaged over voxels in each ROI, were entered into repeated measures ANOVAs, with factors of transition type, transition magnitude, and ROI. For comparisons of pairs of ROIs or transition types, paired two-tailed t-tests were used. Default Bayes Factors (BF_10_) are reported for all F statistics (Faulkenberry & Brennan, 2023) and t statistics (Rouder et al., 2009), with higher numbers (>1) favouring the alternative hypothesis and smaller numbers (<1) favouring the null hypothesis. Whole brain contrasts are shown following smoothing with a 10 mm full-width at half-maximum Gaussian kernel, and with t statistics thresholded by controlling the false discovery rate (FDR) to be <0.05 (Benjamini & Yekutieli, 2001). Visualisation uses MRIcroGL software (https://www.nitrc.org/projects/mricrogl).

## Results

### Co-occurrence of transition types

As expected, for model 1, the correlation between the small and large magnitudes of each transition type was low (r < 0.01), because the time points used to generate them are mutually exclusive (Figure 1A). Location and temporal regressors were more correlated, as transitions in both features often co-occurred. Indeed, the highest correlation was between large temporal and large location transitions (range 0.54 – 0.85 across runs). Figure 1A also shows variance inflation factors for each transition and event boundary regressor for each run of models 1 and 2. For the final run, the sum of the location transition regressors equaled the sum of the temporal transition regressors. Averaging across the other seven runs, all variance inflation factors were < 4.3 (range 1.1 to 7.9 across runs), suggesting an acceptable level of collinearity overall (O’Brien, 2007).

**Figure 1.**
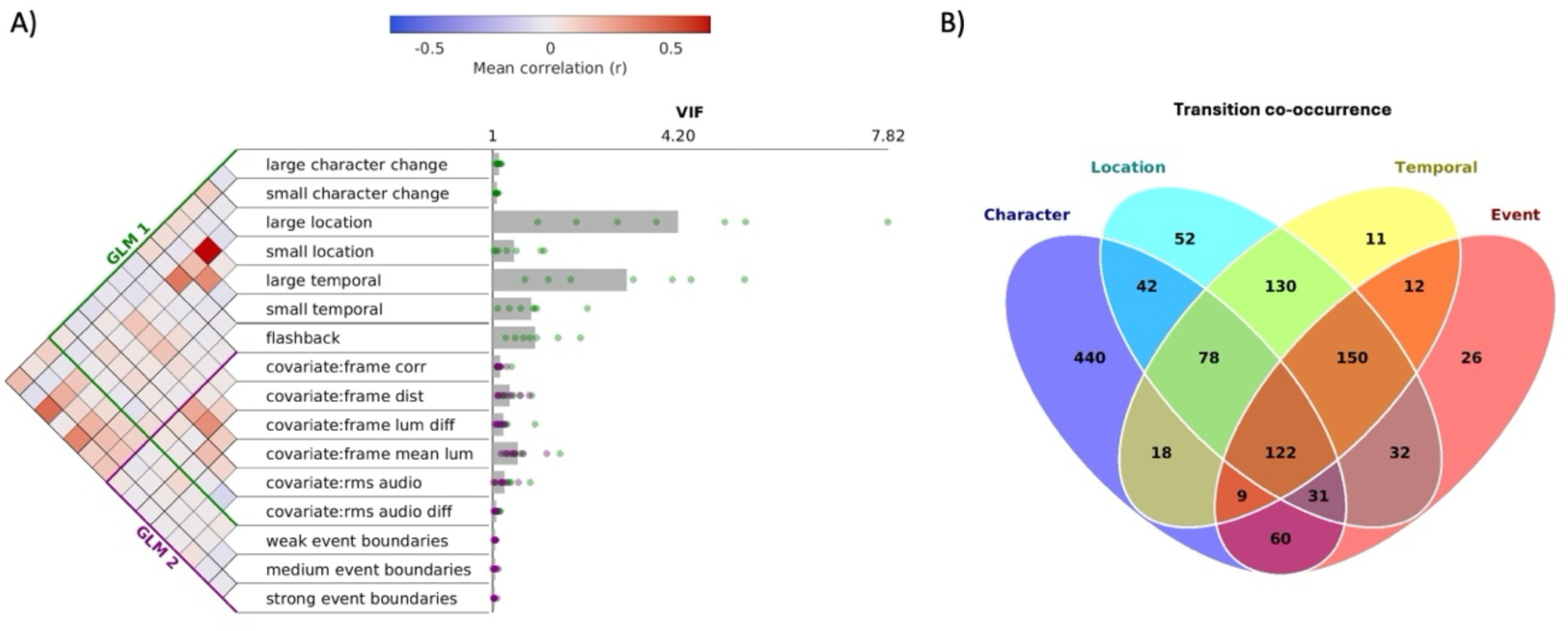
Panel A shows the regressor correlation matrix (Pearson’s r) for the first and second models, with regressors for each type of transition at each magnitude level, plus the audio-visual covariate. The bars show the variance inflation factors (VIF) averaged across runs, with the dots indicating the VIF for individual runs. Panel B shows a Venn diagram indicating the number of occurrences (combined across runs) of each possible combination of single or coincident transition types, as defined in the third model.

We next assessed the frequency with which transitions of different types occurred at the same time or in close temporal proximity (within 2 s). The number of cases for all combinations of transition types, as defined in model 3, are shown as a Venn diagram in Figure 1B, collapsing across the eight runs. While many cases involved multiple transition types occurring together, as expected, there were also many “purer” instances of transitions of each single type.

### DMN subnetworks all respond to narrative transitions, but with different transition type preferences

Focusing on the DMN subnetworks, our first model investigated whether the subnetworks preferentially activate for particular transition types, and whether transition magnitude affects the activation (Figure 2A). A three-way ANOVA with factors of ROI (Core, MTL and dmPFC DMN subnetworks), transition types (location, temporal and character), and transition magnitude (large, small) showed a significant effect of ROI (F_(2,28)_ = 24.95, p < 0.01, BF_10_ = 2.66×10^4^), interaction of ROI and transition type (F_(2,28)_ = 28.12, p < 0.01, BF_10_ = 1.21×10^10^), and interaction of ROI and transition magnitude (F_(2,28)_ = 4.30, p = 0.03, BF_10_ =2.11). Whole-brain activation maps, with DMN subnetworks outlined, are presented in Figure 2B. These maps, and the ANOVA interactions, suggest that the subnetworks exhibit different activation across the various types of transitions. Each ROI is therefore next examined in turn.

**Figure 2.**
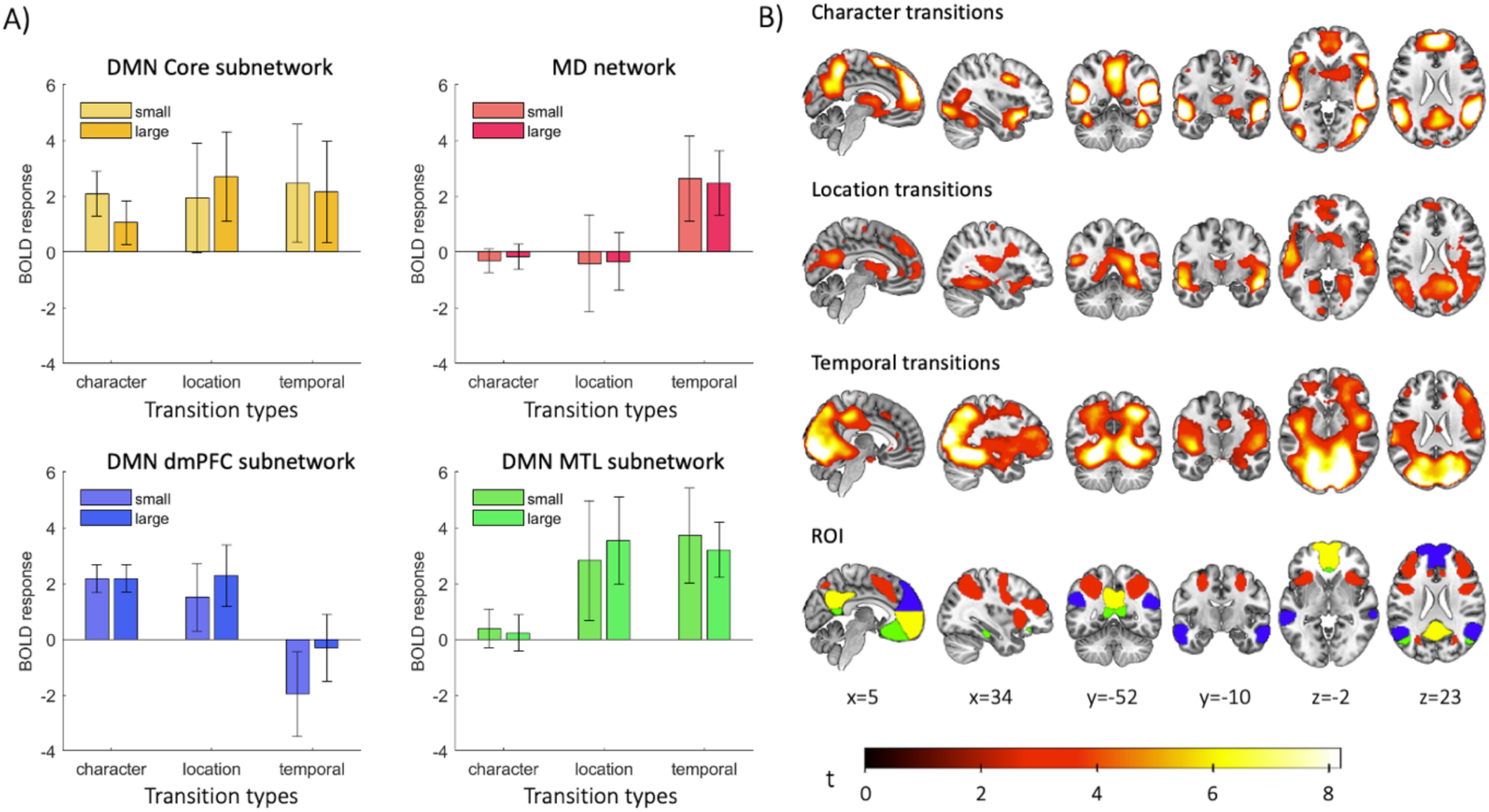
Neural activity during transitions. A. DMN subnetwork and MD activity during each magnitude of character, location and temporal transitions. Error bars indicate between-subject 95% confidence intervals for the contrast of each transition type against baseline activity. B. Whole-brain activity during character, location and temporal transitions, averaged across transition magnitude. Whole-brain activation maps are smoothed with a 10mm full-width at half-maximum Gaussian kernel (FDR-corrected, showing voxels with p <0.05). The DMN subnetwork and MD ROIs are shown in the fourth row, with colours matching the bars in panel A. Slices are labeled with their MNI coordinates (mm).

### The Core DMN responded to all transition types

The Core DMN has been shown to activate during several types of task switches as compared to task repetitions in task-based paradigms (Crittenden et al., 2015; Smith et al., 2018; Zhou et al., 2024a, 2024b), and thus the Core DMN was hypothesised to also activate at transitions in naturalistic stimuli. Indeed, a two-way repeated measures ANOVA found that overall activation at transitions was significantly positive as shown by the intercept (F_(1,14)_ = 33.62, p <0.01, BF_10_ = 297). Further, studies using task-based paradigms demonstrated that with complex task structures, Core DMN activated more during more dissimilar task switches (between stimulus domains) than during more similar task switches (within stimulus domains). Therefore, it was hypothesised that for a larger magnitude of change in each naturalistic stimulus feature, the Core DMN would also exhibit higher activity. However, there was no significant difference in activation between the small and large magnitude levels of narrative transitions (F_(1,14)_ = 0.13, p = 0.73, BF_10_ = 0.37) or between the three transition types (F_(2,28)_ = 0.82, p = 0.42, BF_10_ = 0.15), or interaction (F_(2,28)_ = 0.67, p = 0.46, BF_10_ = 0.13). Thus Core DMN activity demonstrated general sensitivity to transitions in naturalistic stimuli, without strong preference for particular types of transitions. As shown in Figure 2A, out of all the ROIs examined, the Core DMN was the only ROI showing significantly positive activation for all types of switches, averaged across magnitudes (t_14_ > 3.46, p < 3.8 x 10^-3^, BF_10_ >12.45).

### The DMN’s MTL subnetwork preferentially activated at location and temporal transitions

The exploratory analyses of HCP data suggested that the MTL subnetwork responds to scenes and especially to transitions of scenes during movie-watching. In addition, the MTL subnetworks’ proximity to the parahippocampal place area (PPA) may implicate it in representing spatial information (Häusler et al., 2022). Here, as shown in Figure 2, the MTL subnetwork showed preferential activation for location and temporal transitions. A two-way ANOVA with factors of transition types and magnitude revealed a significant effect of transition type (F_(2,28)_ = 21.27, p < 0.01, BF_10_ = 7702), with no significant effect of magnitude (F_(1,14)_ = 0.004, p = 0.98, BF_10_ = 0.35) or interaction (F_(2,28)_ = 0.42, p = 0.59, BF_10_ = 0.11). *Post-hoc* two-tailed t-tests showed that MTL subnetwork activity was significantly higher for both location and temporal transitions than character transitions, averaging over the magnitude levels (t_14_ > 5.13, p < 0.01, BF_10_ >197), with no difference between the two (t_14_= 0.41, p = 0.68, BF_10_ = 0.28). The MTL subnetwork’s activation at character transitions was not significantly above baseline (t_14_ = 1.21, p = 0.25, BF_10_ = 0.49).

### The DMN’s dmPFC subnetwork preferentially activated at character and location transitions

The dmPFC subnetwork also showed distinct activation responses to transitions. A two-way ANOVA with factors of transition types and magnitude found a significant main effect of transition type (F_(2,28)_ = 19.66, p < 0.01, BF_10_ = 4284). In this case, there was also a significant effect of magnitude (F_(1,14)_ = 7.43, p = 0.02, BF_10_ = 3.69), but no significant interaction (F_(2,28)_ =1.53, p = 0.24, BF_10_ = 0.27). Supporting the previous exploratory findings of strong dmPFC activations at character transitions, a *post hoc* two-tailed t-test confirmed significantly positive activations at character transitions (t_14_ = 10.58, p < 0.01, BF_10_ = 2.85 x 10^5^).

Matching the DMN’s Core and MTL subnetworks, the dmPFC subnetwork also had significant activation for location transitions (t_14_ = 4.91, p < 0.01, BF_10_ = 138). However, in contrast to the other three networks, the dmPFC subnetwork deactivated for temporal transitions (t_14_ = 2.17, p < 0.05, BF_10_ =1.59). Extending the results from the significant interaction of ROI and magnitude from the three-way ANOVA, while the other ROIs showed no significant difference between small and large transition magnitudes, the dmPFC subnetwork exhibited significantly higher activation for larger compared to smaller magnitudes of temporal transitions (t_14_ = 2.22, p < 0.05, BF_10_ = 1.73), but not for location (t_14_ = 1.03, p = 0.32, BF_10_ = 0.41) or character transitions (t_14_ = 4.66 x10^-3^, p = 0.99, BF_10_ = 0.26). Additionally, *post hoc* two-tailed t-tests indicated significantly higher dmPFC subnetwork activity at character and location transitions than temporal transitions (t_14_ >3.84, p <0.01, BF_10_ >23.51). Somewhat in contrast from the results for HCP data, there was no significant difference between character and location transitions (t_14_ = 0.78, p = 0.45, BF_10_ = 0.34).

### The MD network responded only to temporal transitions

For the frontoparietal MD ROI, a two-way ANOVA showed a significant effect of transition type (F_(2,28)_ = 22.06, p < 0.01, BF_10_ = 1.01 x 10^4^), with no significant effect of magnitude (F_(1,14)_ = 0.003, p = 0.96, BF_10_ = 0.35) or interaction (F_(2,28)_ = 0.03, p = 0.97, BF_10_ = 0.08). T-tests for each type of transition (averaging over magnitude) showed that there was no significant activation for character or location transitions (t_14_<1.65, p>0.12, BF_10_<0.79), and no significant difference between character and location transitions (t_14_ = 0.32, p = 0.76, BF_10_ = 0.27). However, there was a significantly positive activation for temporal transitions (t_14_ = 5.64, p <0.01, BF_10_ = 445). Two-tailed t-tests confirmed a significantly larger activation for temporal transitions than for location (t_14_ = 4.53, p <0.01, BF_10_ = 73.95) and character transitions (t_14_ = 7.31, p <0.01, BF_10_ = 5.21 x 10^3^).

### Each type of transition preferentially activated specific networks

While the preceding results indicated a specificity of transition types at which each subnetwork preferentially activated, we also compared activity between subnetworks at each transition type. Two-tailed t-tests between each combination of subnetwork pairs per transition type showed that the dmPFC subnetwork responded more to character transitions than did MD and MTL subnetworks (t_14_ >10.91, p < 0.01, BF_10_ > 4.07×10^5^), and somewhat more than core DMN, though with anecdotal BF (t_14_ = 2.21, p = 0.04, BF_10_ = 1.68). The MTL subnetwork responded more to location transitions than did the other DMN and MD ROIs (t_14_>2.67, p <0.01, BF_10_>3.39). For temporal transitions, the MTL subnetwork also exhibited higher activation than the MD or other DMN subnetworks (t_14_>2.17, p <0.05, BF_10_>1.59), and while the MD and Core DMN ROIs showed no difference (t_14_ = 0.41, p = 0.68, BF_10_ = 0.28), both showed higher activation than the dmPFC subnetwork (t_14_>4.81, p <0.01, BF_10_>117).

### All DMN subnetworks responded to subjective event boundaries in a graded manner

Previous research has shown the hippocampus to be sensitive to subjectively-rated event boundaries (Ben-Yakov & Henson, 2018). Given that the nature of event boundaries indicates a transition in the ongoing narrative, we were interested in also assessing whether DMN subnetworks respond to these subjectively-rated event boundaries. Outputs from the second GLM were first averaged across voxels within the ROIs for the hippocampus, DMN subnetworks and frontoparietal MD network, and plotted for each level of boundary salience (Figure 3). A repeated measures ANOVA per ROI was used to test for a mean response to the event boundaries and a main effect of event boundary salience (low, medium, high). First, we reproduced the previously reported finding of hippocampal activation at event boundaries in these data (Ben-Yakov & Henson, 2018). The ANOVA confirmed a significantly positive intercept (F_(1,14)_ =43.93, p <0.01, BF_10_ = 874), and a significant effect of boundary salience in the hippocampal ROI (F_(2,28)_ =18.56, p <0.01, BF_10_ = 2.83 x 10^3^). Next, the DMN subnetworks and MD network were tested with similar one-way ANOVAs. In all cases, results resembled those for the hippocampus: Core subnetwork: a significant intercept (F_(1,14)_ =38.93, p <0.01, BF_10_ = 532), and a significant main effect of salience (F_(2,28)_ =19.93, p <0.01, BF_10_ = 4.73 x 10^3^); MTL: a significant intercept (F_(1,14)_ = 59.21, p <0.01, BF_10_ = 3.17×10^3^), and significant main effect of salience (F_(2,28)_ =17.50, p <0.01, BF_10_ = 1.87 x 10^3^); dmPFC: significant intercept (F_(1,14)_ =32.17, p <0.01, BF_10_ = 251) and significant main effect of salience (F_(2,28)_ =32.46, p <0.01, BF_10_ = 2.41×10^5^); MD: significant intercept (F_(1,14)_ =31.45, p <0.01, BF_10_ = 230) and significant main effect of salience (F_(2,28)_ =16.83, p <0.01, BF_10_ = 1.43 x 10^3^).

**Figure 3.**
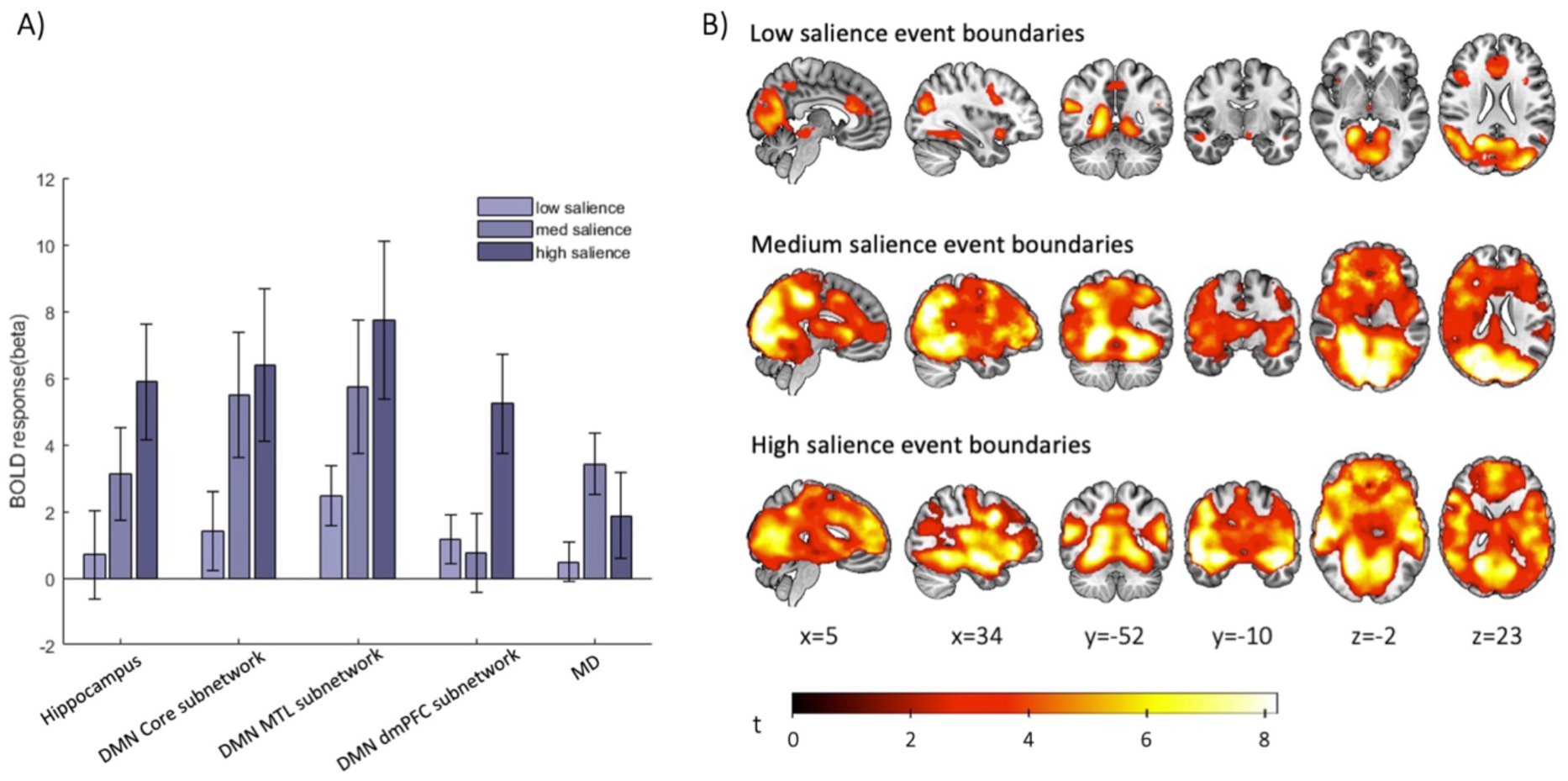
Network activity at subjective event boundaries. A. Hippocampal, DMN subnetworks’, and MD’s BOLD responses at low, medium and high salience levels of subjective event boundaries. Error bars indicate between-subject 95% confidence intervals for the contrast of each transition condition against baseline. B. Whole-brain activity at event boundaries of low, medium and high salience levels. Activation maps are smoothed with a 10mm full-width at half-maximum Gaussian kernel (FDR-corrected, showing voxels with p <0.05).

It is noteworthy that the Core and MTL subnetworks, like the hippocampus, exhibited a monotonic increase with salience. In contrast, MD network activity was significantly higher for medium salience event boundaries, compared to both low and high salience boundaries (t_14_>2.55, p <0.02, BF_10_ >2.84). The whole brain analysis revealed widespread activity for the higher-salience event boundaries. This is consistent with expectations that a subjective event boundary could be identified based on any of various stimulus features that are coded in many different brain regions (Speer, 2009; Zacks 2010). The presence of strong activity in early visual cortex in particular suggests that the confound regressors included in the model may not have been sufficient to explain all low-level perceptual changes associated with event boundaries.

### Consistent response profiles to pure transition types

Our final analysis sought to confirm that distinct transition-response profiles across ROIs were still observed when attempting to isolate pure transitions of each type, and when combining objective transitions and event boundaries in a single model (Figure 4). To investigate the effects of pure transitions, we focused on transitions that occurred without co-occurrence of another transition within a +/− 2 second window. A two-way repeated measures ANOVA with factors ROI (Core, MTL, dmPFC subnetworks, and MD) and transition type (location, character, temporal, and subjective event boundary) confirmed a significant main effect of ROI (F_(3,42)_ = 19.05, p <0.01, BF_10_ = 2.64 × 10^5^), and interaction (F_(9,126)_ = 9.71, p < 0.01, BF_10_ = 4.54 × 10^7^), highlighting that the networks exhibited distinct patterns of activation across different types of pure transitions.

**Figure 4.**
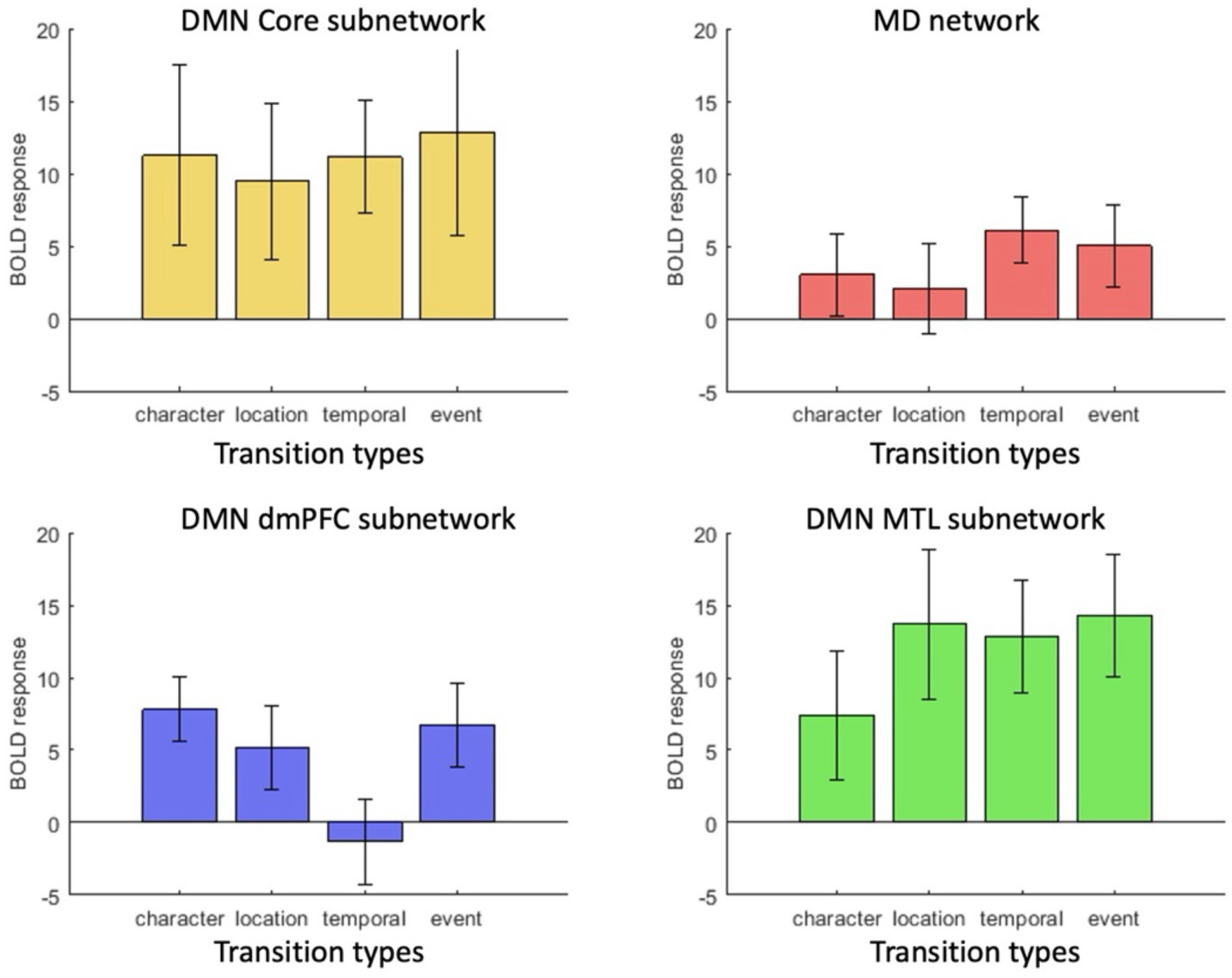
Neural activity during pure transitions. DMN subnetwork and MD activity during independent cases of character, location, temporal and event boundary transitions. Error bars indicate between-subject 95% confidence intervals for the contrast of each transition type against baseline activity.

One-way ANOVAs per ROI generally confirmed the results reported above. In the Core DMN, no significant effect of transition type (F_(2,28)_ = 0.60, p = 0.56, BF_10_ = 0.03) suggested general sensitivity to all types of pure transitions; one-sample t-tests showed significant activation for pure transitions in character (t_14_ = 3.88, p < 0.01, BF_10_ = 25.16), location (t_14_= 3.75, p < 0.01, BF_10_ = 20.16), and time (t_14_= 6.16, p < 0.01, BF_10_ = 980.2), as well as event boundaries (t_14_ = 3.89, p < 0.01, BF_10_ = 25.60).

The MTL subnetwork exhibited a significant main effect of transition type (F_(3,42)_ = 6.01, p < 0.01, BF_10_ = 14.91). Although it was also significantly activated by all pure transitions (character: t_14_ = 3.60, p < 0.01, BF_10_ = 15.50; location: t_14_= 5.70, p < 0.01, BF_10_ = 485.0; temporal: t_14_= 7.08, p < 0.01, BF_10_ = 3.75 x 10^3^; event boundary: t_14_= 7.19, p < 0.01, BF_10_ = 4.38 x 10^3^), pairwise comparisons between transition types indicated that activation at character transitions was significantly lower than at location transitions (t_14_ = –4.26, p < 0.01, BF_10_ = 47.38), temporal transitions (t_14_= –2.59, p = 0.02, BF_10_ = 2.99) and event boundaries (t_14_= –3.54, p < 0.01, BF_10_ = 14.02), with no significant difference between the latter three types (t_14_ < 1.22, p > 0.24, BF_10_ < 0.49).

The dmPFC subnetwork also exhibited a significant main effect of transition type (F_(3,42)_ = 20.42, p < 0.01, BF_10_ = 5.84 x 10^5^), with significant activation for character transitions (t_14_= 7.52, p < 0.01, BF_10_ = 6.94 x 10^3^), location transitions (t_14_ = 3.85, p < 0.01, BF_10_ = 23.94), and event boundaries (t_14_ = 4.99, p < 0.01, BF_10_ = 157.0), but not temporal transitions (t_14_ = – 0.96, p = 0.35, BF_10_ = 0.39); character transitions were associated with significantly greater activation than location transitions (t_14_ = 3.35, p < 0.01, BF_10_ = 10.32) and temporal transitions (t_14_ = 6.99, p < 0.01, BF_10_ = 3.35 x 10^3^), but not event boundaries (t_14_ = 1.06, p = 0.31, BF_10_ = 0.42).

Finally, the MD ROI also exhibited a significant main effect of transition type (F_(3,42)_ = 5.70, p < 0.01, BF_10_ = 11.01), with significant activation for temporal transitions (t_14_ = 5.71, p < 0.01, BF_10_ = 490.1), event boundaries (t_14_ = 3.84, p < 0.01, BF_10_ = 23.20), and marginally for character transitions (t_14_ = 2.28, p = 0.04, BF_10_ = 1.87), but not for location transitions (t_14_ = 1.41, p = 0.18, BF_10_ = 0.60). Temporal transitions produced significantly greater responses than character transitions (t_14_ = 2.71, p = 0.02, BF_10_ = 3.65), and location transitions (t_14_ = 3.26, p < 0.01, BF_10_ = 8.88), but not event boundaries (t_14_ = –1.07, p = 0.301, BF_10_ = 0.43).

To summarize, these results further support the hypothesis that the Core DMN activates to the broadest range of transition types, while the other DMN subnetworks and the MD network show selective sensitivity to specific transition types. The DMN’s dmPFC subnetwork preferred pure character transitions, but also responded to location transitions; the DMN’s MTL subnetwork preferred pure location and temporal transitions, but also responded to character transitions; the MD network preferred pure temporal transitions, with a marginal response to character transitions.

## Discussion

### Differential sensitivity of DMN subnetworks to specific types of narrative transition

The present study presents compelling evidence that the subnetworks of the DMN are sensitive to transitions in different aspects of the narrative context within naturalistic stimuli. The subnetworks’ consistent activations for specific types of transitions suggest a specialised sensitivity to features that contribute to a mental model of an ongoing narrative (Speer et al., 2009; Stawarczyk et al., 2021; Zacks et al., 2010). This is consistent with theories of DMN serving an environmental monitoring function (Buckner et al., 2008; Gilbert et al., 2007; Hahn et al., 2007; Raichle et al., 2001; Shulman et al., 1997), and representing situation models more broadly (Ranganath & Ritchey, 2012; Yazin et al., 2024).

Notably, the Core DMN subnetwork exhibited positive activation across all types of examined transitions, including character, location and temporal transitions. Together with its response to task-based transitions (Crittenden et al., 2015; Smith et al., 2018; Zhou et al., 2024a, 2024b), the Core DMN thus appears to have a rather general sensitivity to cognitive transitions of many kinds. In contrast, the MTL subnetwork showed a clear preference for location and temporal transitions, perhaps partly reflecting proximity to the parahippocampus and associated structures implicated in scene construction and spatial navigation (Burgess et al., 2002; Hassabis & Maguire, 2009), and consistent with a role of similar regions in “mental time travel”(Addis et al., 2007; Buckner & Carroll, 2007). Whereas the MTL subnetwork was least activated by character transitions, the dmPFC subnetwork was strongly activated by character (as well as location) transitions, consistent with roles of the dmPFC, temporal pole and temporoparietal junction in social cognition (Amodio & Frith, 2006; Van Overwalle, 2009), theory of mind (Gallagher & Frith, 2003), and emotion processing (Lettieri et al., 2019; Olson et al., 2007).

Interestingly, while the exploratory analysis of HCP data suggested that the dmPFC subnetwork was more sensitive to character-related than scene-related features, the analyses of *StudyForrest* data suggested a broader involvement of this region, responding robustly to both characters and location transitions (although again with a preference for pure character transitions over pure location transitions). One possibility is that the surrounding context in which transitions occur may influence the network’s engagement. For example, in Forrest Gump, transitions to a known location (e.g. Forrest’s house) may activate associated semantic knowledge (e.g. linked to Forrest himself). In the HCP data, where short film clips were either unfamiliar or presented without the context of the preceding plot, semantic links between characters and locations are likely to have been weaker.

Although there was some correlation between the timing of different transition types (e.g. large temporal transitions often co-occurred with large location transitions), the significant parameter estimates for each transition type in at least two ROIs, plus the significant interactions between each pair of transition types and at least one pair of ROIs, suggests that multi-collinearity did not hamper our ability to distinguish responses to different transition types. This conclusion is reinforced by the final analysis that found similar subnetwork dissociations when isolating more “pure” transition types from compound transitions.

Overall, differentiation between the DMN subnetworks underscores their specialisation in tuning to distinct aspects of contextual information, perhaps suggesting divided functions under an overall monitoring role of the DMN, integrated in the midline Core regions, in line with previous accounts of relative regional specialization (Andrews-Hanna, 2012; Andrews-Hanna et al., 2010, 2014; Axelrod et al., 2017; Wen, Mitchell, et al., 2020).

### Graded response to salience of subjective event boundaries, but not to semantic distance of individual transitions

At subjective event boundaries, we found that all DMN subnetworks responded more strongly to more consistently rated boundaries. This activity profile is comparable to the hippocampus, which was previously argued to play the role of a ‘film-editor’ in segmenting and encoding extended events during continuous movie-watching (Ben-Yakov & Henson, 2018). Whether DMN subnetworks share this role or are involved in cognitive processes upstream or downstream of the encoding remains to be determined.

Interestingly, while results showed a significant effect of salience on all DMN subnetworks’ activity for subjective event boundaries, there was minimal effect of the semantic magnitude of the character, location, and temporal transitions. This could highlight nuanced differences between how the DMN networks process subjective event boundaries versus transitions in the external features that evoke them, or differences in how these types of transition have been defined. High salience event boundaries reflect critical moments in the movie where there is a collective recognition of substantial change, potentially imbuing them with implicit significance that may not be fully contained within the stimulus input. Additionally, the DMN is implicated in predictive coding and anticipating future events based on the past (Addis et al., 2007; Buckner, 2010), and high salience event boundaries may violate these anticipations more significantly than smaller transitions.

### Multiple demand regions responded selectively to temporal transitions and had a unique response profile to subjective event boundaries

While the DMN ROIs each activated for at least two types of transition, the MD network’s activation for only temporal transitions suggest that this network may be recruited for distinct processes required to process and integrate temporal discontinuities. Temporal transitions differ from location and character transitions in that, in addition to recognising, and often integrating, explicit sensory cues that indicate a change, there is a judgement to be made about whether it is a jump forward or backward in the timeline of the story, and by how far.

Further, the temporal transitions in the stimuli could be considered ‘non-natural’ because in real life external jumps in perceived time are rare. Given the MD network’s role in adaptive cognitive control, this activation could relate to involvement in maintaining and updating attention during moments of heightened cognitive demand, similar to its response to suspense in movies (Naci et al., 2014). Extending this observation, MD activation might arise at any disruption in the perceived flow of time, with suspense being a special case when attention is endogenously shifted to a future point in time, in anticipation of an upcoming event.

Recognising a temporal shift requires reorienting oneself to the narrative timeline, and making comparisons to the previous scene and to the anticipation of what should happen if the narrative were continuous. Therefore, MD activation at temporal shifts may reflect particular aspects of processing the change that require working memory, understanding causal relationships and the sequence of events, making predictions, updating the mental model to maintain cohesiveness of the narrative, and reasoning to allow comprehension.

In contrast to the DMN subnetworks’ monotonic increase of activity in response to more salient event boundaries, the MD network was unique in responding most strongly to medium-salience event boundaries. In task-based paradigms, the defining feature of the MD network is greater activation for harder compared to easier tasks (Assem et al., 2020; Fedorenko et al., 2013). It is therefore possible that high-salience event boundaries, with their abundance of visual cues and sharp narrative shifts, are easier to detect, whereas medium-salience boundaries require more implicit judgments to recognize and comprehend, making them cognitively more demanding. Low-salience boundaries, on the other hand, are, by definition, reported by fewer observers, and so may escape the attention of some altogether. An alternative possibility is that DMN subnetworks, such as the dmPFC, peak in activity during high-salience event boundaries and may inhibit the MD network, assuming an intrinsic antagonistic relationship between the networks (Fox et al., 2005; Margulies et al., 2016). The interaction between DMN subnetworks and the MD network—and the specific stimulus or cognitive features that drive their responses to transitions—remains an interesting question for further investigation.

### Limitations

It remains intriguing how far the current results generalise beyond film stimuli, where transitions are entangled with numerous stimulus and narrative properties. For example, a new character appearing might trigger a response because the stimulus is novel, or because it is a person, or because the character is meaningful to the narrative of the story, or because the appearance simply reminded the subject of a familiar acquaintance. In naturalistic stimuli, transition types often co-occur with each other, and are signaled by lower level perceptual properties of the input. The results suggest that we were at least partly able to disentangle responses to the individual transition types examined, although this separation is likely to be incomplete. Similarly, although the inclusion of several visual and auditory covariates suggests that the measured responses reflect more abstract properties of transitions, the presence of widespread activation, including much of occipital cortex, in response to temporal transitions and to high salience event boundaries, suggests that these transitions are associated with other unmodelled visual changes. More controlled experiments are required to test in finer detail which properties of transitions are necessary and sufficient to evoke DMN subnetwork activity.

The magnitude levels chosen here to categorise the transitions are also highly dependent on the available annotations and the type of transition; thus, it was impossible to equate the levels of transition magnitude across types. For example, the larger-magnitude level of character transitions was defined by the first appearance of any character in a scene, which is qualitatively different from the roughly distance-based levels of location and temporal transitions. However, other magnitudes could be used, such as a conceptual distance between characters that are sequentially portrayed on screen, perhaps measured by their familial or social relationship, or the frequency with which the characters appear on screen together.

### Conclusion

The current findings clarify the DMN’s role in externally directed cognition. Studies using task-based paradigms often suggest that DMN is an internally-oriented system (Buckner et al., 2008; Murphy et al., 2018), with roles in mind-wandering (Christoff et al., 2009), self-related thought (Andrews-Hanna et al., 2010; Davey et al., 2016; Gusnard et al., 2001), autobiographical memory retrieval (Addis et al., 2007; Schacter et al., 2007) and mental imagery (Hassabis & Maguire, 2009). However, recent studies have found consistent activation of Core DMN at externally driven task switches (Crittenden et al., 2015; Smith et al., 2018; Zhou et al., 2024a), suggesting also an externally focused cognitive role for the DMN. A review of naturalistic studies also demonstrated that the DMN’s neural responses are modulated by external events unfolding over time (Yeshurun et al., 2021). Instead of a primarily internally-focused model (Andrews-Hanna et al., 2014; Raichle & Snyder, 2007; Smallwood et al., 2011), Yeshurun et al (2021) suggest that the DMN plays the role of a ‘sense-making’ system, interpreting external events in a context-dependent manner. A monitoring role for the DMN has also long been proposed (Buckner et al., 2008; Gilbert et al., 2007; Raichle et al., 2001; Shulman et al., 1997). The current results are consistent with both the sense-making and monitoring perspectives. Together, DMN regions actively track stimulus-based information, responding especially at moments pertaining to a change in the current cognitive model. Specifically, we show that the DMN’s MTL and dmPFC subnetworks have distinct roles in monitoring specific transition types in an attended, external, information-rich narrative, while the Core DMN has a general sensitivity to cognitive transitions of many kinds. We therefore suggest, in line with other contemporary proposals (Stawarczyk et al., 2021; Yazin et al., 2024; Yeshurun et al., 2021), that the Core DMN subnetwork has a special role in integrating external and internal information, processing changes in external conditions to update an internal mental model that, whether at rest, engaged in a task, or absorbed in a movie, structures our current understanding of the world around us.

## Acknowledgments

Ashley X. Zhou, Daniel J. Mitchell and John Duncan were supported by Medical Research Council Intramural Program MC_UU_00030/7. Ashley X. Zhou was supported by a Gates Cambridge Scholarship. We are grateful for open data shared by the *StudyForrest* project (http://www.studyforrest.org), by Ben-Yakov & Henson (2018), and by the Human Connectome Project, WU-Minn Consortium (Principal Investigators: David Van Essen and Kamil Ugurbil; 1U54MH091657) funded by the 16 NIH Institutes and Centers that support the NIH Blueprint for Neuroscience Research; and by the McDonnell Center for Systems Neuroscience at Washington University. For the purpose of open access, the UKRI-funded authors have applied a Creative Commons Attribution (CC BY) licence to any Author Accepted Manuscript arising from this submission.

